# Exposure to childhood maltreatment is associated with changes in sperm small non-coding RNA and DNA methylation profiles

**DOI:** 10.1101/2023.04.27.538231

**Authors:** Jetro J. Tuulari, Matthieu Bourgery, Annukka Ahonen, Ammar Ahmedani, Jo Iversen, Thomas Gade Koefoed, Eeva-Leena Kataja, Linnea Karlsson, Romain Barrès, Hasse Karlsson, Noora Kotaja

## Abstract

**Background:** Childhood maltreatment exposure (CME) increases the risk of adverse long-term health consequences for the exposed individual. Animal studies suggest that CME may also influence the health and behaviour in the next generation offspring through CME-driven epigenetic changes in the paternal germ line. The contribution of paternal early life stress on the health of the next generation in humans is not fully elucidated.

**Methods:** In this study, we measured paternal CME using the Trauma and Distress Scale (TADS) questionnaire and mapped sperm-borne sncRNAs expression by small RNA sequencing (small RNA-seq) and DNA methylation (DNAme) in spermatozoa by reduced-representation bisulfite sequencing (RRBS-seq) in males from the FinnBrain Birth Cohort Study. The study design was a (nested) case-control study, high-TADS (TADS ≥ 39, n = 25 for DNAme and n = 14 for small RNA-seq) and low-TADS (TADS ≤ 10, n = 30 for DNAme and n = 16 for small RNA-seq)). Groups were compared to identify specific epigenetic signatures associated with TADS levels in the spermatozoa of participants.

**Results:** Compared to the control group, high CME was associated with altered sperm sncRNA expression and DNAme profiles. Particularly, we identified several tRNA-derived small RNAs (tsRNAs) and miRNAs with markedly changed levels in males with high CME. DNA methylation analysis identified several genomic regions with differentially methylated CpGs between groups. Notably, we identified two epigenetic marks related to brain development with distinct profiles between CME and controls, the miRNA hsa-mir-34c-5p and differential methylation of the region in proximity of FSCN1.

**Conclusions:** This study provides further evidence that early life stress influences the paternal germ line epigenome and supports a possible contribution in the development of the central nervous system of the next generation.

## INTRODUCTION

Adverse childhood experiences (ACEs) include harms that affect children indirectly through their living environments (e.g. parental conflict, substance abuse, or mental illness) or directly (abuse and neglect). The direct harms are commonly described as childhood maltreatment exposure (CME). CME is highly prevalent, as shown by a recent systematic review and meta analysis that reported a pooled prevalence of ca. 23% in Europe and the U.S. for adults who reported at least one ACE (Hughes et al., 2021). Worldwide, as many as 12% of adults report a history of childhood sexual abuse, 23% of childhood physical abuse, and 36% of emotional abuse (Stoltenborgh, Bakermans-Kranenburg, Alink, & IJzendoorn, 2015). ACEs have numerous adverse consequences for later health, via a range of hormonal, metabolic and immunological pathways (Soares, Rocha, Kelly-Irving, Stringhini, & Fraga, 2021), especially for mental health outcomes (Sara & Lappin, 2017; Stoltenborgh et al., 2015). In addition to affecting health later in life (Waehrer, Miller, Marques, Oh, & Harris, 2020), accumulating evidence indicate that paternal ACEs / CME may also affect the health of the next generation (Dickson et al., 2018; Moog et al., 2018; Roberts et al., 2018; Scorza et al., 2019; Yehuda & Lehrner, 2018).

Animal studies of paternal inheritance induced by early life stress have shown that advert psychological exposures change the epigenetic marks in sperm and the metabolic and behavioural phenotype of the offspring (J Bohacek et al., 2015; Johannes Bohacek & Rassoulzadegan, 2019; K. Gapp et al., 2020; Katharina Gapp et al., 2014; Kretschmer & Gapp, 2022). The commonly recognized epigenetic marks are DNA methylation, histone modifications, and expression of small non-coding RNAs (sncRNA) (Ghai & Kader, 2022; Maamar, Beck, Nilsson, McCarrey, & Skinner, 2022; Nestler, 2016; Wang, Liu, & Sun, 2017). When carried in gametes, these epigenetic marks have the potential to change the early embryonic developmental trajectory and affect offspring phenotype. Numerous studies have identified a link between paternal exposure to a plethora of physical environmental exposures like toxins, cigarette smoking, physical activity, and nutritional stress and changes in sperm epigenome or altered offspring phenotype (Ghai & Kader, 2022; Senaldi & Smith-Raska, 2020). However, the association between early-life psychological stress and epigenetic changes in spermatozoa remains unclear.

To our knowledge, three human studies have established a link between paternal early-life stress and changes in the sperm epigenome have been published to date. In the first, childhood Trauma Questionnaire (CTQ) and Conflict Tactics Scales (CTS) were used to quantify CME in 34 males (17 men exposed to high, 5 men to medium, and 12 men to no childhood abuse), an association was found between DNA methylation and CTQ / composite abuse score (Roberts et al., 2018). In this cohort, twelve DNA regions were differentially methylated between individuals with different childhood abuse (high vs. low), including genes associated with neuronal function (MAPT, CLU), fat cell regulation (PRDM16), and immune function (SDK1) (Roberts et al., 2018). In the second study, CME was quantified using adverse childhood experiences (ACEs) screen; and comparing a group of individuals with the highest ACE score to individuals with the lowest score, a negative correlation was found between levels of multiple miRNAs of the miR-449/34 family and ACE scores (Dickson et al., 2018). In the third study (preprint at the time of writing), there was a longitudinal approach that uses three different age groups. They first report that miR-16 and miR-375 levels are higher in the serum of children exposed to paternal loss and maternal separation (ages 7-12 years), demonstrate similar results in another sample of 18-25-year-olds, and finally used the CTQ to quantify ACEs and find that the same miRNA have lower expression in sperm of adult men exposed to higher CME at ages 21-50 years (Jawaid et al., 2020). Interestingly, they also replicate prior findings by Dickson et al. on lower levels of miR-34 but not miR-449.

In the current study, we measured CME with Trauma and Distress Scale (TADS) questionnaire and quantified sperm sncRNA profiles and DNAme from males that were divided into 2 groups, a control group with low TADS scores (TADS ≤ 10) and a case group with high TADS scores (TADS ≥ 39) to identify between-group differences in CME-associated features in the sperm epigenome. Following the primary analyses, we also performed replication analyses to prior work where possible. This was an exploratory study and there were no a priori hypotheses on the strength or direction of the possible associations between CME and sperm DNA methylation and sncRNA profiles.

## MATERIAL AND METHODS

The study was conducted in accordance with the Declaration of Helsinki and was approved by the Ethics Committee of the Hospital District of Southwest Finland (15.3.2011 §95, ETMK: 31/180/2011). We followed the Strengthening the Reporting of Observational Studies in Epidemiology (https://www.strobe-statement.org/) reporting guideline (case-control studies v4).

### Participants

Participants were recruited at gestational week (gwk) 12 from maternity clinics in Southwest Finland from 2011 to 2015 to take part in the FinnBrain Birth Cohort Study (http://www.finnbrain.fi), which was established to prospectively investigate the effects of early life stress, including prenatal stress exposure, on child brain development and health. The cohort entails 3808 families and included full trios (mother, father, child) on approximately half of the families (L. Karlsson et al., 2018).

The division into case and control groups was based on previously collected data. CME was assessed by questionnaires filled in by the fathers-to-be following the initial recruitment approximately at gwk 14 (2011 - 2015). There was no case - control matching during recruitment although the approach and analyses in the current study are based on case - control design. The new data were collected during visits that took place between 02 / 2019 – 07 / 2021. The recruitment rate for the visits has been 59.6%, which is typical for our cohort study. Until the time of writing, this has been a cross-sectional study.

Altogether 75 males participated in the current study. The final sample size was determined by the available resources. We processed sperm samples from 58 individuals with TADS scores ranging between 0 and 78. 55 samples were analyzed by RRBS-seq to identify DNA methylation patterns, and 30 were analyzed by small RNA-seq (27 samples overlapping). The flowchart of participant selection is presented in Figure 1.

**Figure 1.**
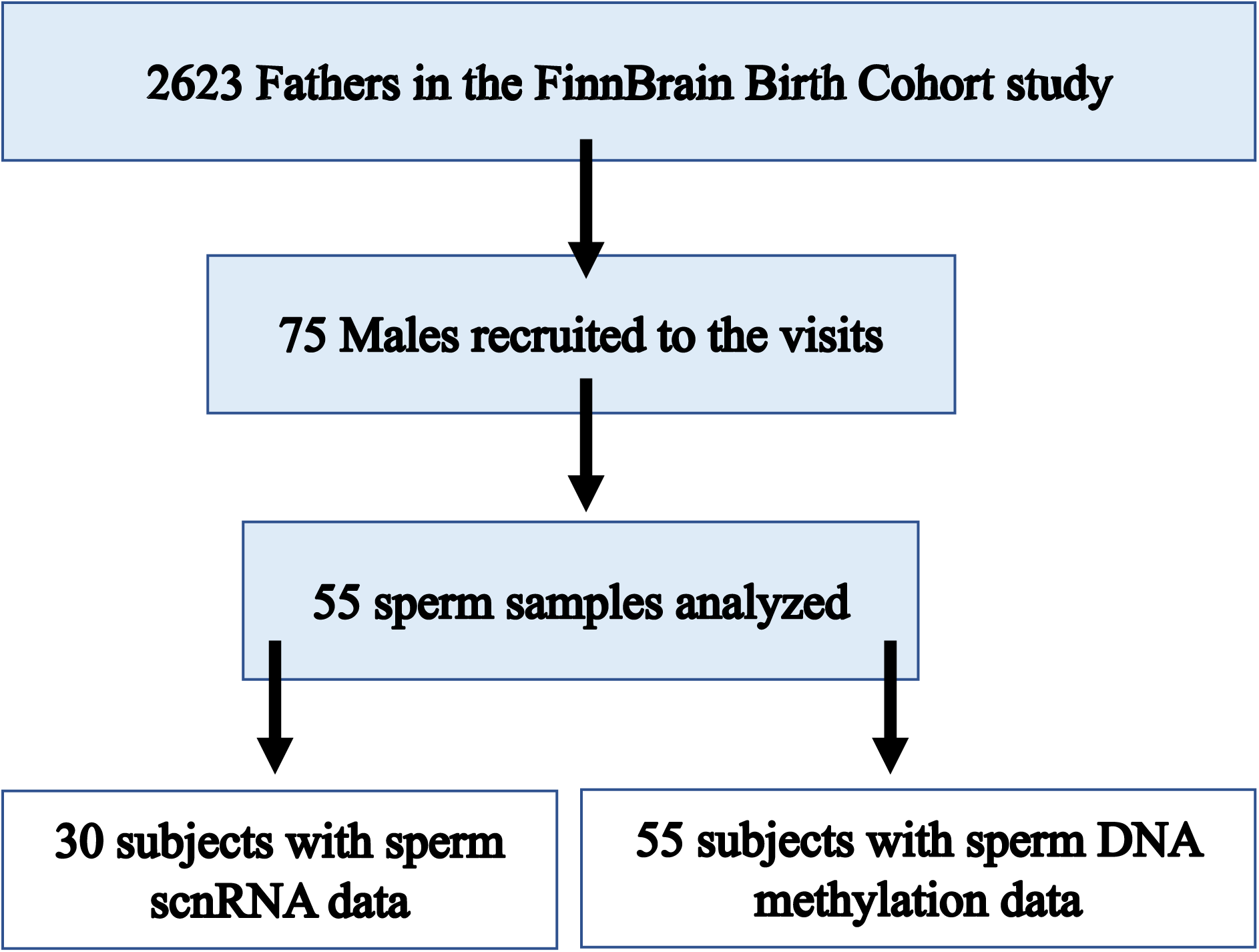
Flowchart of the participant selection for the current study.

The bias assessment is as follows: there was no systematic monitoring for selection bias during recruitment as the primary recruitment for the cohort entailed a population representative sample and was fairly uniform on e.g., for participants’ age range; we strived to quantify the influence of potential confounding factors by including multivariate statistical tests; questionnaire-based measurements are prone to measurement error and TADS questionnaire to recall bias, but these limitations apply to all similar studies; finally we performed the processing of semen samples with utmost care and where possible assured their quality to reduce measurement errors.

### The demographics and questionnaire data

Data were mostly obtained over the prenatal period and data collection was continued postnatally. Information about the parents’ CME was collected using the Trauma and Distress Scale (TADS) at gwk 12 (Salokangas et al., 2016). The TADS comprises five core domains: emotional neglect, emotional abuse, physical neglect, physical abuse, and sexual abuse. In this study, we calculated the cumulative exposure to early life stress events of the infants’ fathers and mothers by the age of 18 years (direct sum scores). Of note, the TADS scores have two possible derived values so that one can use direct sum scores or sums of factor scores (Salokangas et al., 2016). Here we used the direct TADS sum scores as the primary variable to perform case-control divisions and statistical testing, but we have used the factor sum scores in our prior article (H. Karlsson et al., 2020) The direct and factor sum scores were highly correlated (in our sample Spearman’s rho = 0.954, p < 0.001).

Depressive symptoms were assessed by implementing the Edinburgh Postnatal Depression Scale (EPDS) (Cox, Holden, & Sagovsky, 1987). This 10-item questionnaire is scaled from 0 to 30 points with a bigger score denoting increased symptom severity. Anxiety symptoms were quantified with the anxiety subscale of Symptom Checklist 90 (SCL-90) (Holi, Sammallahti, & Aalberg, 1998). The SCL-90 anxiety subscale consists of 10 items with a total score range of 0 to 40 with larger scores denoting increased symptom severity. Alcohol use was quantified with a structured questionnaire (Danielsson et al., 2022), and the derived variable we chose was mean alcohol use per week (1 unit = 12 g of alcohol). Smoking was quantified with a yes / no binary variable. BMI was obtained from height and weight measurements performed during the research visits that are described below.

### The study visits

Participants that matched the case-control criteria (control group TADS ≤ 10; and case group TADS ≥ 39) were invited to a separate study visit. During the visit fathers filled in questionnaires for smoking and alcohol use habits, depressive symptoms (EPDS), and anxiety symptoms (SCL- 90). They also gave additional biological samples (not reported here). Weight and height were measured for defining body mass index (BMI), and waist circumference was measured following standard procedures. Sperm samples were collected before or at the visit.

### The sperm sample collection

Participants were allowed two options for delivering the semen samples following 2-7-day abstinence from ejaculation: either collecting the sample at home and delivering it to the research site at the start of the visit or giving the sample during the visit. The semen was collected by masturbation. Obtained samples were incubated at +37°C for 5-30 min for liquefaction, and spermatozoa were purified by centrifuging through 50% Puresperm (Nidacon) solution at 400xg for 15 min. The sperm pellet was subsequently washed with mild somatic cell lysis buffer (0.01% SDS, 0.005% Triton X-100) to eliminate remaining somatic cell contamination. The purity of the samples was assessed by light microscopic analysis. The total number of spermatozoa before purification ranged between 40 and 800 million (only one high TADS sample had less than 40 million spermatozoa), and after purification, all samples were pure with only very minor somatic cell contamination (Supplementary Figure 1A). Sperm purification was conducted promptly within the same day and purified samples were frozen for storage.

### Small RNA sequencing analysis

Total RNA was extracted from 14 high-TADS (TADS ≥ 39) and 16 low-TADS (TADS ≤ 10) samples (10 million spermatozoa per sample) by TRIzol LS (Invitrogen) containing tris(2- carboxyethyl)phosphine (TCEP, Sigma-Aldrich) as a reducing agent to enhance sperm nucleus lysis, and precipitated with isopropanol in the presence of 2 µl of GlycoBlue (15 mg/ml, Invitrogen). After DNaseI treatment (Sigma-Aldrich) without heat inactivation, RNA was re extracted with Trizol LS to remove DNaseI. The quality of the RNA sample was analyzed by Bioanalyzer (Agilent RNA 6000 Pico Kit). Bioanalyzer analysis validated the lack of somatic cell contamination, as demonstrated by the absence of ribosomal RNA peaks and low RIN values (2-3) (Supplementary Figure 1B). The RNA yield from 10 million spermatozoa was 10-200 ng.

Libraries for small RNA-seq were prepared using NEB Next® Multiplex Small RNA kit (New England Biolabs), and the libraries were sequenced by NovaSeq 6000 system (Illumina).

The quality of the reads was assessed using FastQC (v0.11.9) (http://www.bioinformatics.babraham.ac.uk/projects/fastqc/). The adapters and bad quality reads were trimmed off using cutadapt (v3.5) (Martin, 2011). SPORTS1.1 (Shi, Ko, Sanders, Chen, & Zhou, 2018) was used to align and map the 15-45 nucleotides long reads first to the human genome (hg38), then subsequently to ribosomal RNAs (rRNAdb), miRNAs (miRBase v22), YRNA mapping and transfer RNAs (tRNA) (GtRNAdb v2.0) using software default settings. Sequences mapped to tRNAs were annotated as 5’ end-, 3’ end or 3’ CCA end-derived according to their location on the parental tRNA sequence. All sperm samples showed typical size distribution of averaged sncRNA reads, including a peak at 21-23 nt for miRNAs, a peak at 31-32 nt for YRNAs, a peak at 31-32 nt for tRNA-derived small RNAs (tsRNAs), and a peak at 31- 33 nt for PIWI-interacting RNAs (piRNAs) (Supplementary Figure 1 C, D). All sequences mapping to rRNAs, miRNAs, and tRNAs were extracted from SPORTS1.1 as a text output file. For piRNA analysis, reads were mapped to piRNA clusters (Ha et al., 2014) in the Human genome (UCSC: Hg38) using HISAT2 (v2.1.0) (Kim, Paggi, Park, Bennett, & Salzberg, 2019), and assigned and counted using featureCounts (v2.0.0) (Liao, Smyth, & Shi, 2014) against reference gtf files. Altogether, we identified unfiltered normalized reads mapping to a total of 838 miRNAs, 266 tsRNAs, 6195 piRNA genomic clusters, 8 rRNA genes (12S-rRNA, 16S-rRNA, 18S-rRNA, 28S- rRNA, 45S-rRNA, 5.8S-rRNA, 5S-rRNA, other-rRNA) and 4 YRNA genes (YRNA1, 3, 4 and 5). Raw read counts were filtered to a minimum of 10 total counts across all 35 samples and normalized.

Methodological differences to prior work: In contrast to the earlier study by Dickson et al. that used miRNA microarrays for initial screening of differentially expressed miRNAs (Dickson et al., 2018), we chose a genome-wide high throughput approach and used small RNA-seq to identify differentially expressed miRNAs in men exposed to high CME compared to men with low CME.

### RRBS library preparation

Sperm DNA was extracted from ∼ 10 million spermatozoa/sample by lysing cells with RLT+ buffer (Qiagen) with 1% Beta-mercaptoethanol for 15 min., and then passing the lysate through AllPrep DNA Mini Spin Columns (Qiagen) and proceeding with manufacturer’s instructions. The quality control was performed by measuring the concentration of DNA samples on a Qubit fluorometer using Qubit dsDNA High Sensitivity Assay Kit (Thermo Fisher). The reduced-representation bisulfite sequencing (RRBS-seq) was performed by the Single-Cell Omics platform at the Novo Nordisk Foundation Center for Basic Metabolic Research, University of Copenhagen. RRBS libraries were constructed from 100 ng genomic DNA using the Ovation® RRBS Methyl-Seq library preparation kit (Tecan) according to the manufacturer’s instructions. Final libraries were quantified by Qubit (Thermo Fisher) and quality checked on Bioanalyzer (Agilent). Pooled libraries were subjected to either a 101-bp single-end sequencing on a NovaSeq 6000 platform (Illumina) or a 76-bp single-end sequencing on a NextSeq 500 platform (Illumina). A total of 5.5 billion reads were generated.

Methodological differences to prior work: The previous study by Roberts et al. (Roberts et al., 2018) identified differentially methylated sperm DNA using methylation BeadChips, but we chose a sequencing-based approach and used RRBS. Of note, these methods have been shown to cover different CpG loci (Carmona et al., 2017).

### DNA methylation analysis

FASTQ files were first generated using bcl2fastq (v. 2.20.0), and subsequently processed by a python script to add the corresponding UMI reads to the FASTQ read headers. As described in The Analysis Guide for NuGEN Ovation RRBS Methyl-Seq (https://github.com/nugentechnologies/NuMetRRBS), the FASTQ files were then trimmed using Trim Galore! (v. 0.6.4) with default settings and further processed using the trimRRBSdiversityAdaptCustomers.py script to remove diversity adapter sequences. Methylation coverage was then extracted using Bismark by aligning reads to the GRCh38 assembly. Here, the deduplicate_bismark step was performed using the --barcode option to deduplicate reads based on the UMIs. Furthermore, the --ignore 3 parameter was used in the methylation extraction step to disregard restriction enzyme sites.

The resulting methylation coverage files were then analysed using the BiSeq R package (v. 1.36.0). In brief, all 55 sperm DNA samples were split into 2 groups, low-TADS (TADS < 10, n = 30) and high-TADS (TADS > 40, n = 25). As described in the BiSeq user guide (https://bioconductor.org/packages/release/bioc/vignettes/BiSeq/inst/doc/BiSeq.pdf), clusters of CpGs along the genome were defined according to the following criteria: a minimum of 20 CpG sites with coverage of >80 % in both TADS-groups and a maximum distance of 100 bp. The methylation level was then smoothed within each CpG cluster, weighted by the coverage of the individual CpGs. To reduce bias due to unusually high coverage, the coverage for each CpG was limited to the 90% quantile.

### Statistical analyses

In line with prior work, we used between-group comparisons to identify features associated with CME in the sperm epigenome, comparing those with low to those with high CME. These discovery analyses were followed by multivariate analyses that included potential confounders in partial correlation models. Third, we performed replication analyses of Dickson et al. 2018. The multivariate and replication analyses were performed with JASP 0.16.3 (https://jasp-stats.org/). Most variables reported in Table 1 had non-normal distributions and as this applied to the majority of sperm epigenetic variables as well, we thus used non-parametric statistics in multivariate and replication analyses.

**Table 1.**
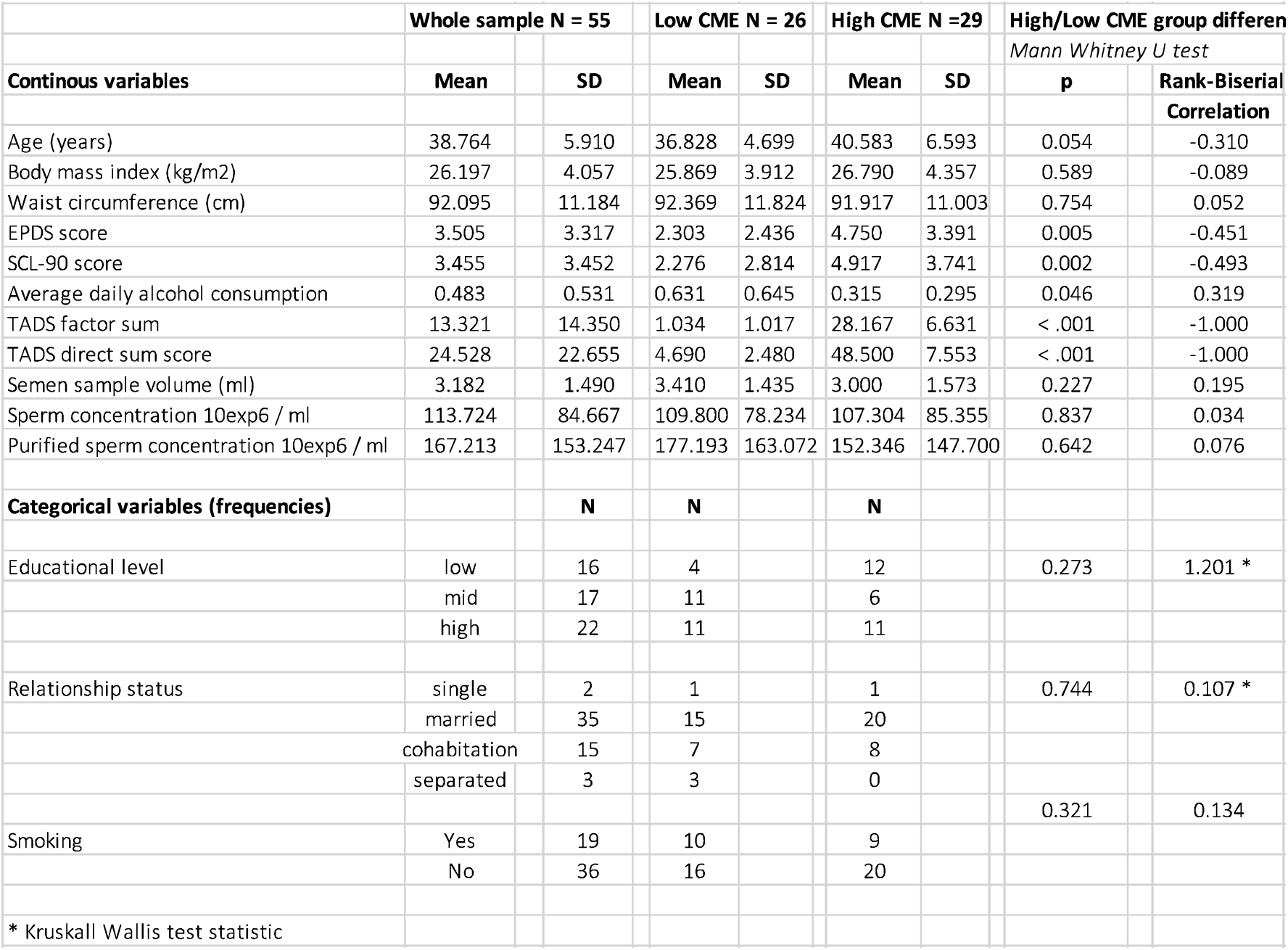
Descriptive statistics of demographics

| Table 1. Descriptive statistics of demographics |  |  |  |  |  |  |  |  |
| --- | --- | --- | --- | --- | --- | --- | --- | --- |
|  | Whole sample N = 55 |  | Low CME N = 26 |  | High CME N =29 |  | High/Low CME group differences |  |
|  |  |  |  |  |  |  | Mann Whitney U test |  |
| Continous variables | Mean | SD | Mean | SD | Mean | SD | p | Rank-Biserial Correlation |
| Age (years) | 38.764 | 5.910 | 36.828 | 4.699 | 40.583 | 6.593 | 0.054 | -0.310 |
| Body mass index (kg/m2) | 26.197 | 4.057 | 25.869 | 3.912 | 26.790 | 4.357 | 0.589 | -0.089 |
| Waist circumference (cm) | 92.095 | 11.184 | 92.369 | 11.824 | 91.917 | 11.003 | 0.754 | 0.052 |
| EPDS score | 3.505 | 3.317 | 2.303 | 2.436 | 4.750 | 3.391 | 0.005 | -0.451 |
| SCL-90 score | 3.455 | 3.452 | 2.276 | 2.814 | 4.917 | 3.741 | 0.002 | -0.493 |
| Average daily alcohol consumption | 0.483 | 0.531 | 0.631 | 0.645 | 0.315 | 0.295 | 0.046 | 0.319 |
| TADS factor sum | 13.321 | 14.350 | 1.034 | 1.017 | 28.167 | 6.631 | < .001 | -1.000 |
| TADS direct sum score | 24.528 | 22.655 | 4.690 | 2.480 | 48.500 | 7.553 | < .001 | -1.000 |
| Semen sample volume (ml) | 3.182 | 1.490 | 3.410 | 1.435 | 3.000 | 1.573 | 0.227 | 0.195 |
| Sperm concentration 10exp6 / ml | 113.724 | 84.667 | 109.800 | 78.234 | 107.304 | 85.355 | 0.837 | 0.034 |
| Purified sperm concentration 10exp6 / ml | 167.213 | 153.247 | 177.193 | 163.072 | 152.346 | 147.700 | 0.642 | 0.076 |
| Categorical variables (frequencies) |  | N | N |  | N |  |  |  |
| Educational level | low | 16 | 4 |  | 12 |  | 0.273 | 1.201 * |
|  | mid | 17 | 11 |  | 6 |  |  |  |
|  | high | 22 | 11 |  | 11 |  |  |  |
| Relationship status | single | 2 | 1 |  | 1 |  | 0.744 | 0.107 * |
|  | married | 35 | 15 |  | 20 |  |  |  |
|  | cohabitation | 15 | 7 |  | 8 |  |  |  |
|  | separated | 3 | 3 |  | 0 |  |  |  |
|  |  |  |  |  |  |  | 0.321 | 0.134 |
| Smoking | Yes | 19 | 10 |  | 9 |  |  |  |
|  | No | 36 | 16 |  | 20 |  |  |  |
| * Kruskal Wallis test statistic |  |  |  |  |  |  |  |  |

### Discovery analyses (case vs. control comparisons)

Differential expression analysis of sncRNAs was performed using DESeq2 (v1.36.0) which is based on a negative binomial generalized linear model (Love, Huber, & Anders, 2014). RsRNA and YRNAs were analyzed using a Wilcoxon-rank exact test. All 30 samples were split into 2 groups, low-TADS (TADS ≤ 10, n = 16) and high-TADS (TADS ≥ 39, n = 14). Low-TADS group was defined as the control group.

For DNA methylation, 55 sperm DNA samples were split into 2 groups, low-TADS (TADS ≤ 10, n = 30) and high-TADS (TADS ≥ 39, n = 25). The statistical tests employed by BiSeq, based on a beta-binomial generalized linear model (GLM), were used to detect CpG clusters and individual CpG sites that were differentially methylated between the two TADS-groups in a false discovery rate (FDR)-controlled manner. Furthermore, the significant CpG sites were used to construct differentially methylated regions (DMRs), which marks regions of significant hyper or hypomethylation within each TADS-group. Finally, CpGs were annotated according to their genomic context, i.e. present in a promoter (within 3kb of the TSS), exon, intron, or as a distal intergenic CpG.

Multiple comparison correction for both scnRNA and DNAme analyses was carried out using Benjamini and Hochberg for sncRNA and DNAme analyses (Benjamini & Hochberg, 1995).

### Multivariate analyses (case vs. control comparisons with covariates)

We performed three different multivariate analyses with partial correlations testing for the association between low vs high CME: A) controlling for semen sample volume and sperm concentration; B) controlling for health characteristics including age, BMI, smoking (yes / no), average alcohol use per day, as well as depressive and anxiety symptoms at the time of the sperm sample collection; C) controlling for covariates in A + B.

### Replication analyses of prior work (miRNA)

We also performed replication analyses of Dickson et al. 2018 for the sncRNA data by exploring the associations between miRNA that their work implicated important and ACE exposure. For this, we looked at between-group differences (low vs. high ACE exposure; and Spearman correlations between CME (TADS scores) and miRNA expression levels for hsa-miR-34c-5p and hsa-miR-449a. We chose to use the TADS factor scores here as the variable had a “more normal” distribution (correlation to TADS sum score Spearman rho = 0.954). We extended these analyses also to partial correlation models where we controlled for age and BMI at measurement, and smoking (yes / no).

## RESULTS

We analyzed the association of the CME as quantified by TADS scores with the two indices of sperm epigenome: the abundance of sncRNAs and levels of DNA methylation. The study enrolment is presented as a flowchart (Figure 1), demographics are reported in Table 1.

### Childhood maltreatment exposure is associated with modified sperm sncRNA profile

The analysis of small RNA-seq data from 14 high-TADS and 16 low-TADS sperm samples showed that the abundance of five analysed classes of sncRNAs (miRNAs, tsRNAs, rsRNAs, YRNAs and piRNA clusters) was generally similar between high-TADS and low-TADS groups (Figure 2A). The vast majority of sncRNA reads originated from rsRNAs and YRNAs (Figure 2A). The reads derived from YRNA and rRNA genes were similarly distributed in high and low-TADS sperm samples, with some differences in the relative number of reads derived from 12S-rsRNA, 16S- rsRNA, 5.8-rsRNA, RNY1 and RNY4 between high and low-TADS samples (Supplementary Figure 1E).

**Figure 2:**
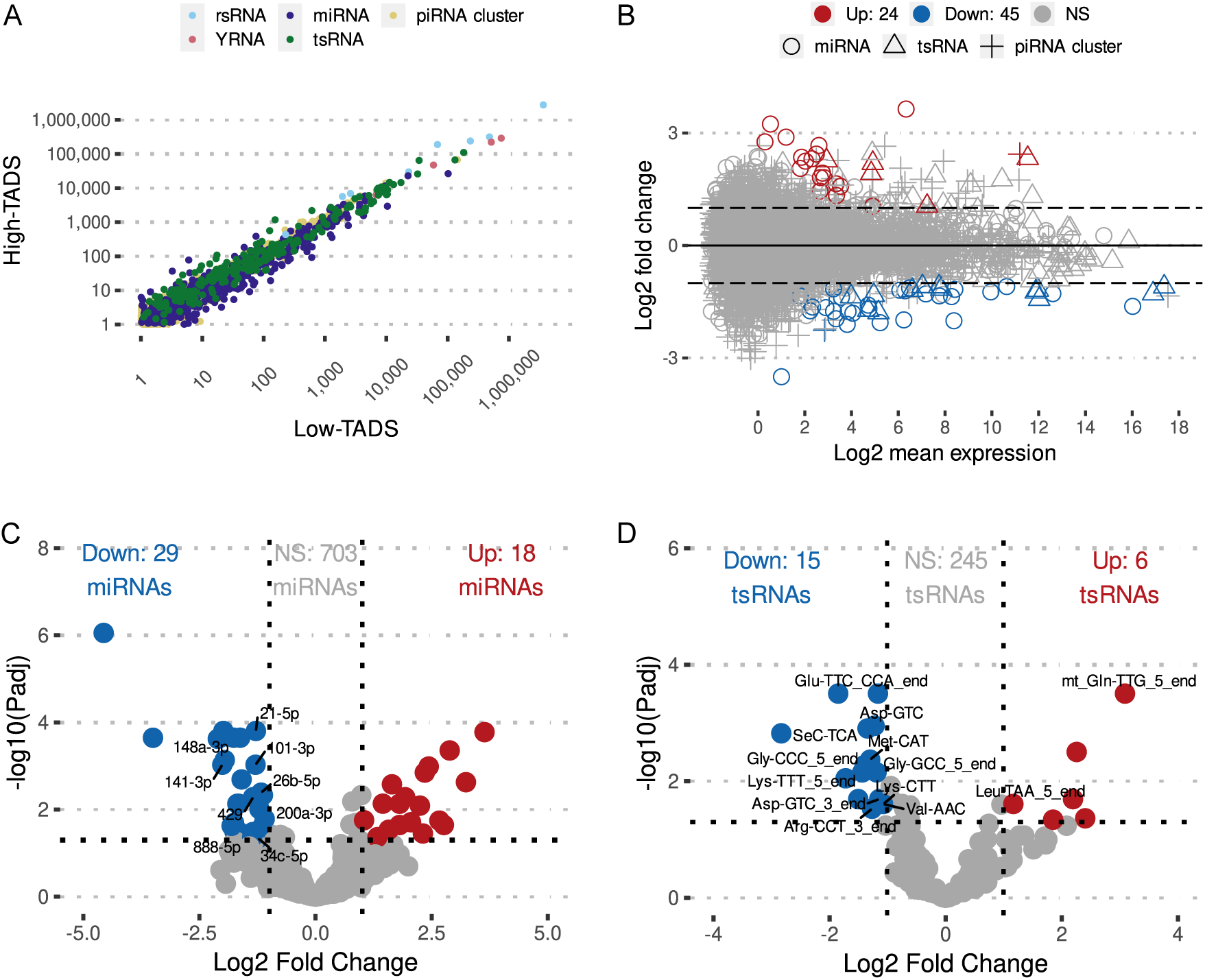
Differential levels of sperm sncRNAs in high-TADS vs low-TADS sperm samples. (A) sncRNA expression between high and low-TADS samples from unfiltered data for low read counts. One dot represents the median of normalized counts of all samples belonging to either high or low-TADS group. (B) MA plot displays the relationship between the log2 fold change in high-TADS vs low-TADS samples and log2 mean expression for all miRNAs, tsRNAs and piRNA clusters. (C) Volcano plots shows differential expression of miRNAs in high vs low-TADS samples. (D) Volcano plots shows differential expression of tsRNAs in high vs low-TADS samples. Blue dots in (C) and (D) visualize sncRNAs with a log2 fold change < - 1.0 (Padj < 0.05). Red dots visualize the sncRNAs with a log2 fold change > 1.0 (Padj < 0.05). Only sncRNAs having baseMean (average of the normalized count values, dividing by size) > 100 are annotated.

The differential expression analysis of individual miRNAs, tsRNAs, and piRNA clusters revealed significant differences in high-TADS vs. low-TADS sperm (Figure 2). A total of 29 miRNAs, 15 tsRNAs, and 3 piRNA clusters were downregulated in high-TADS sperm compared to low-TADS sperm (log2FC < -1.0 and P_adj_ < 0.05), while 18 miRNAs, 6 tsRNAs, and 1 piRNA cluster were upregulated (log2FC > 1.0 and P_adj_ < 0.05) (Figure 2 B-D). The expression levels of three tsRNAs and four miRNAs showing the most significant changes (log2FC > 2, and P_adj_ < 0.01) and the number of normalized counts among all samples was above the threshold “baseMean > 10”. Interestingly, hsa-miR-34c-5p, which was earlier shown to be downregulated in sperm samples of individuals with high ACE scores (Dickson et al., 2018; Jawaid et al., 2020), was also downregulated in our discovery analysis.

### Childhood maltreatment exposure is moderately associated with modified sperm CpG methylation

To investigate the possible association between early life stress experience and sperm DNA methylation, we performed RRBS-seq to analyze the methylation levels of CpG-rich regions. We found only very modest differences between high and low-TADS sperm samples. A total of 45 differentially methylated CpGs were unveiled with FDR-corrected p < 0.15. The most significant differences (FDR p < 0.15, abs (meth.diff) > 0.1) were observed for eight CpGs within chromosome 7 (Figure 3). The identified CpGs were all in the same cluster located in the intergenic region, and the closest gene to this differentially methylated region (DMR) is the actin-bundling protein Fascin 1 (FSCN1), which has been shown to control the migration of neuroblasts (Sonego et al., 2013). While this DMR is distant from the FSCN1 gene and not located within a mapped enhancer region, it may affect embryonic neurogenesis after fertilization.

**Figure 3.**
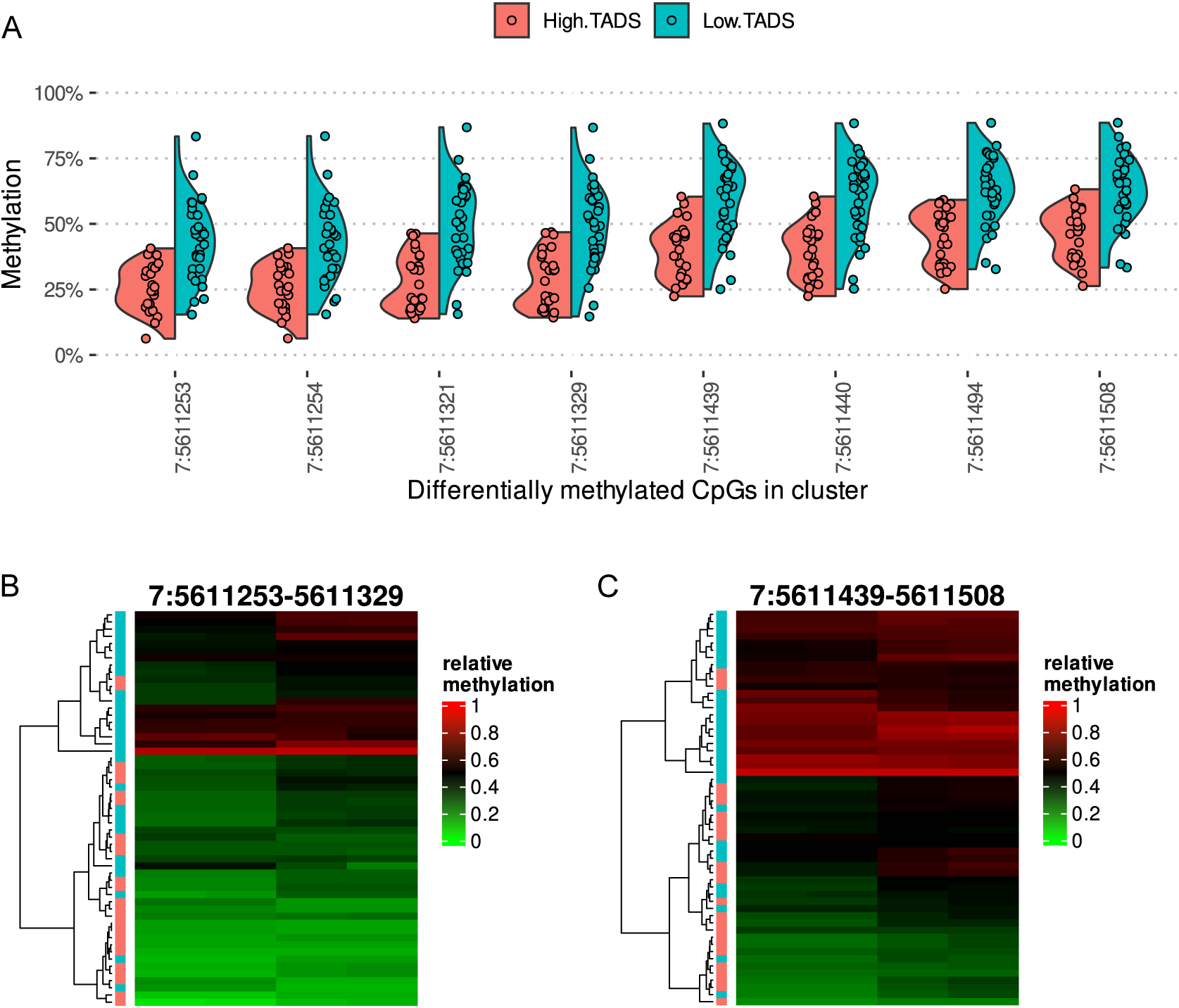
Differential DNA methylation on chromosome 7 in high-TADS vs low-TADS sperm samples. (A) Violin plots shows differentially methylated CpGs in high and low-TADS samples. Only CpGs with FDR threshold <= 0.15 and abs(meth.diff) > 0.1 are shown. (B) Heatmap shows four CpGs from cluster 7_519 from locations 7:5611253, 7:5611254, 7:5611321, and 7:5611329. (C) Heatmap shows four CpGs from cluster 7_519 from locations 7:5611439, 7:5611440, 7:5611494, and 7:5611508. Each row in (B) and (C) represents one sample, with blue cells corresponding to low-TADS and red cells to high-TADS samples.

### CME associations are generally robust potential confounders in multivariate analyses

We identified the following potentially relevant covariates and confounders, based on prior literature, that were also measured in the current study: sperm sample volume, sperm concentration, age, BMI, smoking, alcohol use, depressive symptoms, and anxiety symptoms. The multivariate statistical analyses showed that the associations that were implicated in the discovery analyses were very robust to include covariates. Most of the associations remained statistically significant even after controlling for all potential confounders.

### Expression of hsa-miR-34c-5p is robustly associated with CME

Some associations between TADS scores and miRNA expression levels were previously identified (Dickson et al. 2018). To investigate the robustness of our discoveries, we investigated to which extent our sncRNA expression results overlapped with the Dickson et al. study. Interestingly, we also found a negative association between ACE score and hsa-miR-34c- 5p with a similar effect size but found that hsa-miR-449a levels were not different between groups (Figure 4; replicating Figure 2 plots of Dickson et al.). However, we replicated the tight association between the relative expression of the two miRNAs (Figure 4). We then performed partial correlation analyses for the depicted associations and controlled for age and BMI at measurement, and smoking (yes / no). The associations were as follows: hsa-miR-34c-5p vs. TADS score (r = -0.528, *p* = 0.001); hsa-miR-449a vs. TADS score (r = -0.185, *p* = 0.339). Of note, the association of hsa-miR-34c-5p vs. TADS score was slightly weaker when controlling for all covariates (r = -0.420, *p* = 0.073). Second, we also performed corresponding between-group comparisons. We replicated the lower expression levels in ACE exposed group for hsa-miR-34c- 5p (W = 140, p = 0.017, rank biserial correlation [rbc] = 0.538) and that there were no differences in expression levels of hsa-miR-152-3p and hsa-miR-375-3p. Our analyses thus show that sperm expression of hsa-miR-34c-5p is robustly negatively associated with CME.

**Figure 4.**
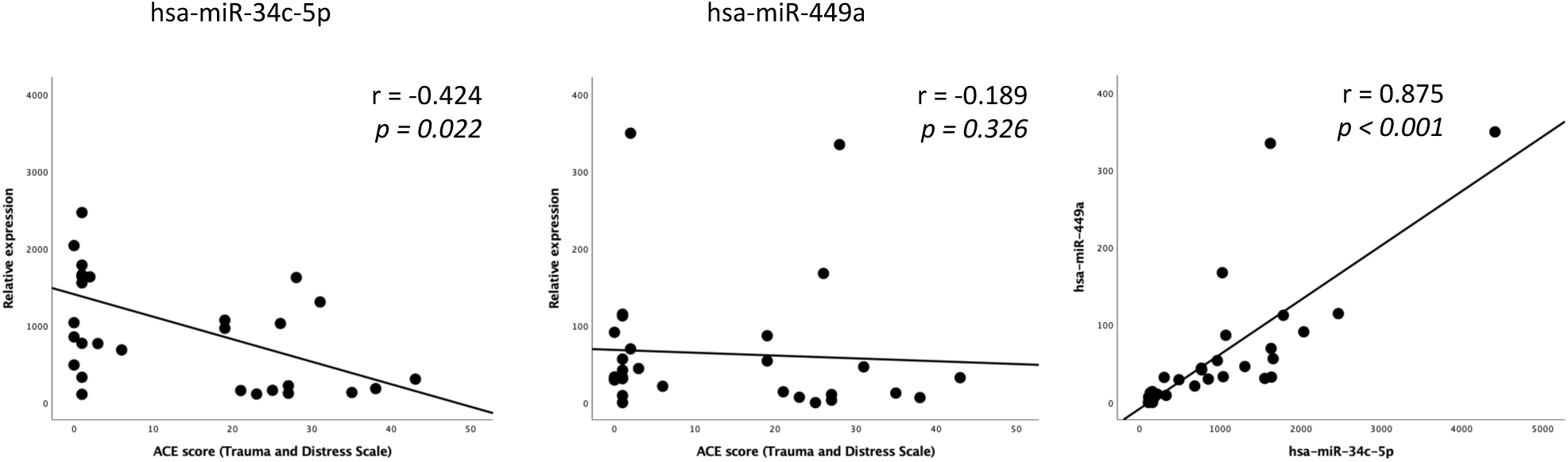
The associations between TADS scores and miRNA expression levels that were identified in prior work (Dickson et al. 2018) [replicating plots in their Figure 2]. We replicated the negative association between ACE score and hsa-miR-34c-5p, but did not replicate the findings on hsa-miR-449a.

## DISCUSSION

Here, we report that CME is associated with multiple epigenetic marks in sperm. We identified differential expression of numerous sncRNAs and a DMR located in the proximity of the *FSCN1* gene in males with high CME. We found an interesting overlap with previous reports for miRNAs, particularly miR-34c-5p, although, most of our results were distinct from prior reports. This may not be surprising given that the methods for quantifying the epigenome and early life stress exposures, i.e., the exposure and outcomes varied between the studies. Our study provides novel insight into the relationship between early life psychological stress and altered sperm epigenome, with possible implications for development in the next generation offspring.

### Questionnaires of early life stressors

Early life adversity and stress are usually assessed retrospectively, and such measures are inherently prone to recall bias and they are unable to capture exposures for the youngest ages, e.g. under the age of three years. It is clear, however, that these questionnaires capture useful information about the cumulative stressors that have been present during childhood. Attesting to this ACEs have been associated with multiple adverse health outcomes (Bellis et al., 2019; Hughes et al., 2017, 2021; Sara & Lappin, 2017).

We measured early life stress with TADS questionnaire that captures five dimensions of neglect and abuse CME that are also part of the most widely adopted CTQ. The other frequently adopted option is to use questionnaires that quantify ACEs. For instance, the ACE Study Questionnaire includes yes / no answers to 10 questions involving participants’ experiences at home until the age of 18. Five of the questions probe CME: physical abuse, verbal abuse, sexual abuse, physical neglect, and emotional neglect which are also the main features of CTQ and TADS questionnaires. The other five questions probe adversity of family members (that has likely a negative influence on the exposed individual): a caretaker with alcoholism or alcohol abuse, experiencing domestic violence, caretaker incarceration, a family member diagnosed with a mental illness, and the loss of a caretaker through divorce, death, or abandonment (Dickson et al. 2018). In summary, there are many options for quantifying ACEs and CME (Thabrew, Sylva, & Romans, 2011), and while most of them probe the five main types of childhood maltreatment, there is very little knowledge on similarities and differences between the questionnaires (Thabrew et al., 2011).

### Epigenetic measures

The stability of the sperm epigenome is not well known. Indeed, human studies frequently use cross-sectional data, which creates obvious limitations for quantifying measurement error (this limitation applies to the current study as well). Roberts et al. (Roberts et al., 2018) report an intra-class correlation coefficient (ICC) between replicate sperm samples taken ca. 3 months apart for the DNAme profiles that were associated with abuse exposure so that ICC values were higher than 0.7 for 90% of implicated sites. Dickson et al. did not collect replicate samples (Dickson et al. 2018). The field would benefit from larger-scale studies that describe the “normative” sncRNA and DNAme profiles in the sperm and describe, which components are stable and which ones are more dynamic across months and longer-term e.g., over several years. Commonly used measures of sperm cell epigenetics, small RNAs and DNA methylation (DNAme) patterns, are modifiable through lifestyle factors, health, and environmental exposures (Ghai & Kader, 2022). Correspondingly, if ACEs cause epigenetic programming in germ line cells that are relevant to intergenerational inheritance, most of them should be relatively stable following the exposure and they would not increase or decrease in time, which is in contrast to many other studied exposures such as cigarette smoking, exercise, acute stress, diet, and obesity.

### Possible links from sperm epigenome to offspring brain development

Although epigenetic mechanisms following fertilization related to DNAme and sncRNAs are likely essential in typical development and may be closely intertwined (Ghai & Kader, 2022; Kretschmer & Gapp, 2022), the most intriguing and robust evidence supports sncRNAs in conveying epigenetic inheritance (Kretschmer & Gapp, 2022; Ostermeier, Miller, Huntriss, Diamond, & Krawetz, 2004; Sendler et al., 2013). Rodent studies have shown transgenerational inheritance of paternal ACEs via changes in miRNA profiles (Johannes Bohacek & Rassoulzadegan, 2019; K. Gapp et al., 2020; Katharina Gapp et al., 2014; Luo, Tan, Li, & Ding, 2022; Rodgers, Morgan, Leu, & Bale, 2015). Similar effects have been reproduced without paternal exposure by injecting the implicated miRNA into zygotes (Dickson et al., 2018), (Katharina Gapp et al., 2014; Rodgers et al., 2015).

Prior work implicated that ACEs were negatively associated with the abundance of miRNAs 449/34 in sperm (Dickson et al., 2018; Jawaid et al., 2020). Importantly, these miRNAs are unexpressed in oocytes but transmitted to them upon fertilization (Dickson et al., 2018; Yuan et al., 2015), and are key regulators of brain development (Grad et al., 2022; Jauhari, Singh, Singh, Parmar, & Yadav, 2018; Mollinari et al., 2015; Wu et al., 2014), including fetal human brain development (Rao et al., 2016; Venø et al., 2017), and possibly also later in development (Morgunova & Flores, 2021). Our analyses provided partial replication for prior work by implicating a negative association between ACEs and miR-34 (Figure 4). As a novel and interesting finding, we identified 21 tsRNAs which had differential abundance in high vs low CME participants. In addition to miRNAs, tsRNAs have also been linked to epigenetic inheritance and could be a biomarker for CME in line with miRNAs (Park, Ahn, Shin, Kim, & Chang, 2020).

Prior studies and the current study identified sperm epigenetic features that could potentially have effects on offspring brain development, which ties in with our recent neuroimaging studies that link paternal CME with offspring neonate brain structure (H. Karlsson et al., 2020; Tuulari, Kataja, Karlsson, & Karlsson, 2022; Tuulari et al., 2023). Several studies, including from our group, have identified that lifestyle factors remodel DNA methylation near genes controlling the development of the brain in human sperm (Donkin et al., 2016; Ingerslev et al., 2018), supporting that genomic regions involved in brain development are hotspots of epigenetic variation in response to environmental stress. While intergenerational effects carried by the sperm epigenome in humans have not been definitively demonstrated, an altered gametic epigenome after CME may influence the development of the central nervous system and modulate the behaviour of the next generation offspring.

### Sperm epigenome as a biomarker of childhood ACEs / CME

Epigenetics is a nascent field with typically small sample sizes that are related to high costs and the need for special infrastructure. This is not too different from the early stages of fields such as genetics and neuroimaging. Within this context, collecting very large data sets is not always feasible. Future studies would benefit from including replicate sperm samples to at least part of the participants to quantify the stable / repeatable elements in both early life stress exposure and control groups. This could potentially be used to identify the most relevant measures of interest for later analyses and greatly decrease the need for multiple comparisons correction. Early life stressors are challenging to quantify reliably, and it may well be that the effects of the exposures are different for different ages, for instance before and after puberty. Prospective cohorts that have detailed information over childhood and measures of early life stressors, including ACEs and CME, could share light on the matter by including sperm data collection e.g., during early adulthood.

Roberts et al. were able to estimate a parsimonious epigenetic marker for childhood abuse using an elastic net model (penalized regression), which identified three DNAme probes that predicted high vs. no childhood abuse in 71% of participants (Roberts et al., 2018). Such findings are very promising and may lead to parsimonious predictive models in the future. Future mechanistic studies should advance our understanding of how DNAme are affected by the underlying genome, and how sncRNAs and DNAme interact in causal pathways following fertilization. Dickson et al. combined data from human and animal models and were able to show that the effects of their CSI stress paradigm implicate similar sncRNA profiles in mice and that these effects are transmitted to embryos and can thus have inter/transgenerational effects (Dickson et al., 2018). Studies that focus on RNA are much better able to measure such epigenetic signatures since DNA methylation undergoes erasure and reestablishment following fertilization, which makes studying the immediate post-fertilization effects challenging (Ghai & Kader, 2022).

### Practical implications

Intergenerational transmission of well-being, health and disease is an important research topic with many implications for health care and societies. It has been postulated that a key component of ACEs, CME is the single most important preventable risk factor for future mental health (Bellis et al., 2019; Hughes et al., 2017, 2021; Sara & Lappin, 2017). CME has also been shown to have effects on health outcomes even when genetic confounding is taken into account (Baldwin et al., 2023). Total annual costs attributable to ACEs were estimated to be US$581 billion in Europe and $748 billion in North America (Bellis et al., 2019; Hughes et al., 2021). Over 75% of these costs arose in individuals with two or more ACEs. Elucidating the mechanisms of intergenerational epigenetic inheritance in humans should be of high priority. Finding interventions to intergenerational effects that pass from one generation to the other could potentially spare future generations from the exposures of their ancestors.

### Strengths and limitations

We used the largest sample size to date for identifying epigenetic marks relates to CME, but the sample size is still modest and larger sample sizes are needed in the future. Measurement error and test-retest reliability of sperm epigenome were not quantified, which could be done with repeated measurements of sperm epigenome. CME measures were obtained retrospectively and are prone to recall bias, but CME measures are widely used and provide the only tangible way of assessing childhood experiences. All participants were Scandinavian / Caucasian, which makes the source population homogenous but necessitates the inclusion of more ethnically diverse populations in the future.

## Conclusions

In the current study, multiple sncRNAs were implicated to have associations with CME. Importantly, for the majority of the sncRNAs, the associations were robust to statistically controlling for semen, sperm, and health-related factors. This is promising for future studies since these findings imply that earlier exposures, even from as far as childhood, can leave an epigenetic mark in sperm cells that is not sensitive to later lifestyle or health.

Taken together, there are clear implications that childhood maltreatment exposure is associated with sperm DNAme and sncRNA profiles. Our results had very little overlap with prior reports, limited to miR-34, which is at least partly due to the variable methodology used for defining the exposure and the epigenetic analyses. Still, this study adds to the evidence that early life stress has influences on adult sperm epigenome. It remains crucial to assess, whether this has ramifications for offspring outcomes in humans.

## Supporting information

Suppelementary Figure 1

## Acknowledgements

We thank all men who participated to the study visits. In addition, we also thank our research nurses for their valuable contribution to the data collection. We want to thank Elina Louramo, Anna Puisto, Tiina Peromaa and Johanna Järvi for technical assistance in organizing the semen collection and performing the sperm purification.

## Author contributions

- Jetro J. Tuulari lead the manuscript writing, participated to the conceptualization of the study, performed statistical analyses.
- Matthieu Bourgery preprocessed the sncRNA data, had a key role in data visualization, performed statistical analyses with major input to the discovery analyses.
- Annukka Ahonen prepared and helped in implementation of the data collection,
- Ammar Ahmedani optimized the sperm RNA extraction protocol and prepared the sperm RNA samples for sequencing.
- Jo Iversen, preprocessed the DNAme data.
- Thomas Gade Koefoed preprocessed the DNAme data.
- Eeva-Leena Kataja provided expertise on interpretation of the CME data and results.
- Linnea Karlsson conceptualized the study and build the infrastructure of the FinnBrain Birth Cohort.
- Romain Barrès provided supervised the DNAme analyses and provided expertise on sperm epigenetics.
- Hasse Karlsson conceptualized the study and build the infrastructure of the FinnBrain Birth Cohort.
- Noora Kotaja conceptualized the study and build the infrastructure for the sperm sample storage and analyses, and supervised the sncRNA analyses.

All authors participated in writing and critically revising the manuscript and accepted it in its final form.

