## Supplementary material for "Exposure to childhood maltreatment is associated with changes in sperm small non-coding RNA and DNA methylation profiles": Suppelementary Figure 1

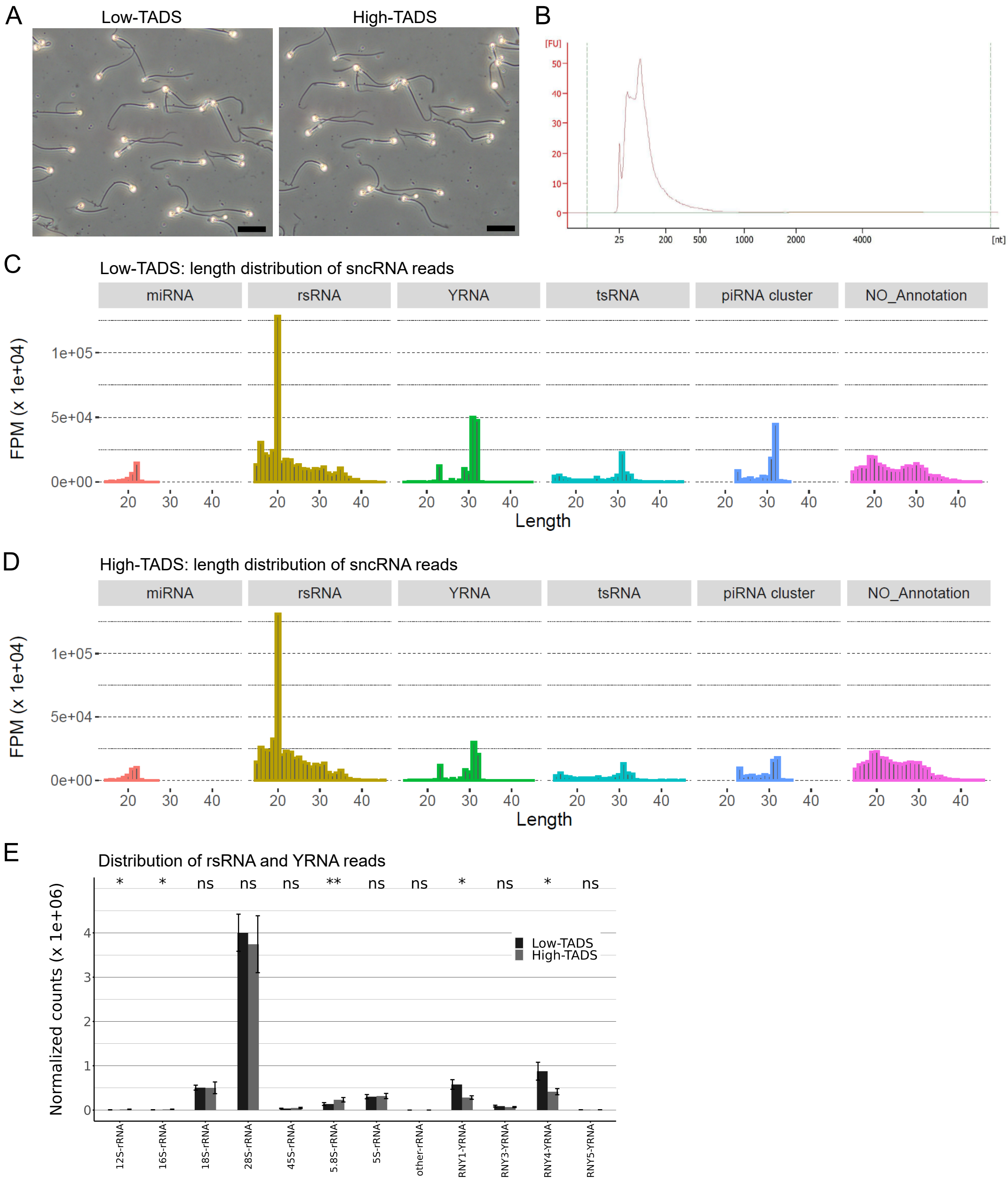


**Supplementary Figure 1. Quality control of sperm samples. (A)** Phase contrast images of representative unstained sperm samples from low-TADS and high-TADS individuals. Scale bars: 20 µm. **(B)** Representative bioanalyzer profile of sperm total RNA sample (low-TADS). **(C, D)** Size distribution of sperm sncRNA reads, representative examples of low-TADS (C) and high-TADS (D) samples are shown. FPM: average fragments per million. Size of sncRNA reads is indicated in nucleotides on the x-axis (Length). **(E)** Distribution of rsRNA and YRNA reads in low- and high-TADS sperm samples. The bars represent the means ± standard error of the normalized reads (FPM, fragment per million) mapping to different types of rsRNAs and YRNAs in low-TADS (n = 16) and high-TADS (n = 14) samples. Wilcoxon-rank exact test indicated significant differences in the abundance of 12S-rsRNA (W=162, p=0.03827), 16S-rsRNA (W=170, p=0.0152), and 5.8-rsRNA (W=177, p=0.005991), as well as for RNY1 (W=58, p=0.02454) and RNY4 (W=57, p=0.02184).
